# A 3D transcriptomics atlas of the mouse olfactory mucosa

**DOI:** 10.1101/2021.06.16.448475

**Authors:** Mayra L. Ruiz Tejada Segura, Eman Abou Moussa, Elisa Garabello, Thiago S. Nakahara, Melanie Makhlouf, Lisa S. Mathew, Filippo Valle, Susie S.Y. Huang, Joel D. Mainland, Michele Caselle, Matteo Osella, Stephan Lorenz, Johannes Reisert, Darren W. Logan, Bettina Malnic, Antonio Scialdone, Luis R. Saraiva

## Abstract

The sense of smell helps us navigate the environment, but its molecular architecture and underlying logic remain unknown. The spatial location of odorant receptor genes (*Olfrs*) in the nose is widely thought to be independent of the structural diversity of the odorants they detect. Using spatial transcriptomics, we created a genome-wide 3D atlas of the mouse olfactory mucosa (OM). Topographic maps of genes differentially expressed in space reveal that both *Olfrs* and non-*Olfrs* are distributed in a continuous and overlapping fashion over five broad zones in the OM. The spatial locations of *Olfrs* correlate with the mucus solubility of the odorants they recognize, providing direct evidence for the chromatographic theory of olfaction. This resource resolved the molecular architecture of the mouse OM, and will inform future studies on mechanisms underlying *Olfr* gene choice, axonal pathfinding, patterning of the nervous system, and basic logic for the peripheral representation of smell.

## INTRODUCTION

The functional logic underlying the topographic organization of primary receptor neurons and their receptive fields is well-known for all sensory systems, except olfaction (*1*). The mammalian nose is constantly flooded with odorant cocktails. Powered by a sniff, air enters the nasal cavity, until it reaches the olfactory mucosa (OM). There, myriad odorants activate odorant receptors (*Olfrs*) present in the cilia of olfactory sensory neurons (OSNs), triggering a complex cascade of events that culminate in the brain and result in odor perception (*1, 2*). In mice, most mature OSNs express a single allele of one out of ~1100 Olfr genes (*Olfrs*) (*3–6*). Olfrs employ a combinatorial strategy to detect odorants, which maximizes their detection capacity (*4, 7*). OSNs expressing the same *Olfr* share similar odorant response profiles (*4, 7*), and drive their axons to the same glomeruli in the olfactory bulb (*8–10*). Thus, *Olfrs* define functional units in the olfactory system, and function as genetic markers to discriminate between different mature OSN subtypes (*6, 11*).

Another remarkable feature of the OSN subtypes is their spatial distribution in the OM. Early studies postulated that OSNs expressing different *Olfrs* are spatially segregated into four broad areas within the OM called ‘zones’ (*12, 13*), but recent work has suggested up to nine overlapping zones (*14–16*). The functional relevance of these zones is not fully understood. These studies, combined, have sampled only ~10% of the intact mouse *Olfr* repertoire. We do not currently understand the full complexity of the OM and, most importantly, lack an unbiased and quantitative definition of zones. In effect, the exact number of zones, their anatomical boundaries, their molecular identity and their potential functional relevance are yet to be determined.

One hypothesis is that the topographic distribution of *Olfrs*/OSN subtypes evolved because it plays a key role in the process of *Olfr* choice in mature OSNs and/or in OSN axon guidance (*17, 18*). An alternative hypothesis is that the spatial organization of *Olfrs*/OSN subtypes is tuned to maximize the detection and discrimination of odorants in the peripheral olfactory system (*12*). Interestingly, the receptive fields of mouse OSNs vary with their spatial location (*19*), which in some cases correlate with the patterns of odorant sorption in the mouse OM – this association was proposed as the ‘*chromatographic hypothesis*’ decades before the discovery of the *Olfrs* (*20*), and later rebranded as the ‘*sorption hypothesis*’ in olfaction (*21, 22*). While some studies lend support to these hypotheses (reviewed in (*23*)), others question their validity (*24, 25*). Thus, the logic underlying the peripheral representation of smell in the peripheral olfactory system still remains unknown, and it is subject of great controversy (*23, 26*).

Spatial transcriptomics, which combines spatial information with high-throughput gene expression profiling, has expanded our knowledge of complex tissues, organs, or even entire organisms (*27–31*). In this study, we employed a spatial transcriptomics approach to create a 3D map of gene expression of the mouse nose, and we combined it with single-cell RNA-seq, machine learning and chemoinformatics to resolve its molecular architecture and shed light into the anatomical logic of smell.

## RESULTS

### A high-resolution spatial transcriptomic map of the mouse olfactory mucosa

We adapted the RNA-seq tomography (Tomo-seq) method (*30*) to create a spatially resolved genome-wide transcriptional atlas of the mouse nose. We obtained cryosections (35 μm) collected along the dorsal-ventral (DV), anterior-posterior (AP), and lateral-medial-lateral (LML) axes (n=3 per axis) of the OM (Figure 1A), and performed RNA-seq on individual cryosections (see STAR Methods). After quality control (Figure S1A-D; Table S1; STAR Methods), we computationally refined the alignment of the cryosection along each axis, and we observed a high correlation between biological replicates (Figure 1B; STAR Methods). Hence, we combined the three replicates into a single series of spatial data including 54, 60 and 56 positions along the DV, AP and LML axis, respectively (Figure 1C; STAR Methods). On average, we detected >18,000 genes per axis, representing a total of 19,249 unique genes for all axes combined (Figure 1D). Molecular markers for all canonical cell types known to populate the mouse OM were detected among all axes (Figure 1E), and were expressed at the expected levels (*6*).

**Figure 1.**
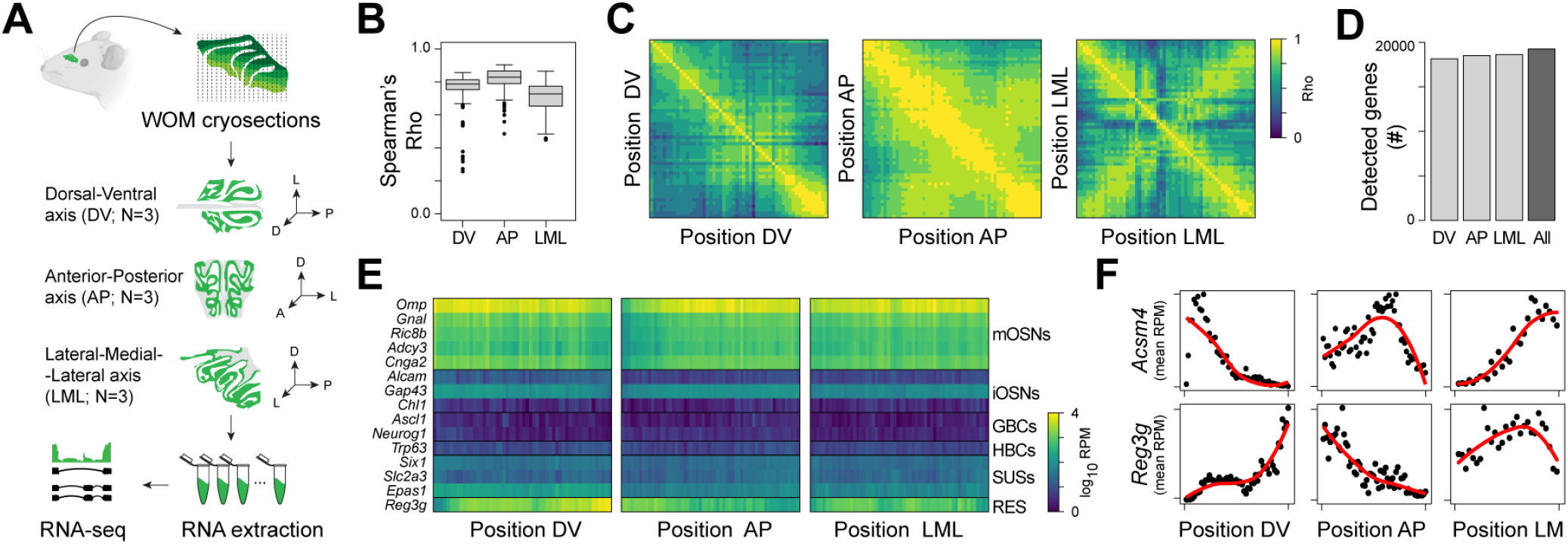
Application of TOMO-seq to mouse OM. (A) Experimental design. TOMO-seq was performed on 9 tissue samples, from which 3 were sliced along the Dorsal-Ventral axis, 3 along the Posterior-Anterior axis and 3 along the Lateral-Medial-Lateral axis. (B) Boxplots showing the distributions of Spearman’s correlation coefficients (Rho) between replicates in each axis. (C) Heatmaps showing Spearman’s correlation between gene expression patterns at different positions along the three axes. (D) Number of detected genes along each axis separately or across the whole dataset. Genes were considered as detected when they had at least one normalized count in at least 10% of the samples from one axis. (E) Heatmaps of log10 normalized expression (after combining the three replicates per axis) of OM canonical markers along the 3 axes (RPM = reads per million; mOSNs = mature Olfactory Sensory Neurons; iOSNs = immature Olfactory Sensory Neurons; GBCs = Globose Basal Cells; HBCs = Horizontal Basal Cells; SUSs = Sustentacular cells; RES = Respiratory epithelium cells). (F) Normalized expression of canonical OM spatial marker genes along the 3 axes. Red line showing fits with local polynomial models.

Next, we verified the presence of a spatial signal with the Moran’s I (*32*) (Figure S1E), whose value is significantly higher than 0 for the data along all axes (p < 2.2×10^−16^ for all axes), indicating that nearby sections have more similar patterns of gene expression than expected by chance. Given the left/right symmetry along the LML axis (Figure 1C), the data was centered and averaged on the two sides (see STAR Methods) – henceforth the LML axis will be presented and referred to as the lateral-medial (LM) axis. We could reproduce the expression patterns in the OM for known spatial markers, including the dorsomedial markers *Acsm4* and *Nqo1* (*33, 34*), and the complementary ventrolateral markers *Ncam2* and *Reg3g* (*35, 36*) (Figures 1F and S1F).

Together, these results show that RNA tomography is both a sensitive and reliable method to examine gene expression patterns in the mouse OM.

### Spatial differential gene expression analysis identifies cell type-specific expression patterns and functional hotspots in the OM

Over the last 3 decades, multiple genes with spatially segregated expression patterns across the OM have been identified. The majority of these genes are expressed in mature OSNs, and include genes encoding chemosensory receptors, transcription factors, adhesion molecules, and many molecules involved in the downstream signaling cascade of chemosensory receptor activation (*6, 13, 14, 16, 33, 34, 36–48*). A smaller number of zonally expressed genes (including xenobiotic compounds metabolizing enzymes, chemokines and transcription factors) were found to be expressed in sustentacular cells, globose basal cells, olfactory ensheathing cells, Bowman’s gland cells, and respiratory epithelial cells (*36, 42, 47–53*). Despite this progress, our knowledge on what genes display true zonal expression patterns, and what cell types they are primarily expressed in is still very limited.

The generation of axis-specific gene expression maps allowed us to explore the relationship between the different axis-specific DEGs and also to discover which cell types they are expressed in, by combining our data with a previously published single-cell RNA-seq dataset (*54*). To identify axis-specific differentially expressed genes (hereafter referred to as spatial DEGs), we first filtered out lowly expressed genes, then binarized the expression levels at each position according to whether they were higher or lower than their median expression, and applied the Ljung-Box test to the autocorrelation function calculated on the binarized expression values (Figure S2A; STAR Methods). After correcting for multiple testing, we obtained a total of 12,303 unique DEGs for the 3 axes combined (FDR<0.01; Figure 2A). Of these, the AP axis showed the highest number of DEGs (10,855), followed by the DV axis (3,658), and the LM (1,318).

**Figure 2.**
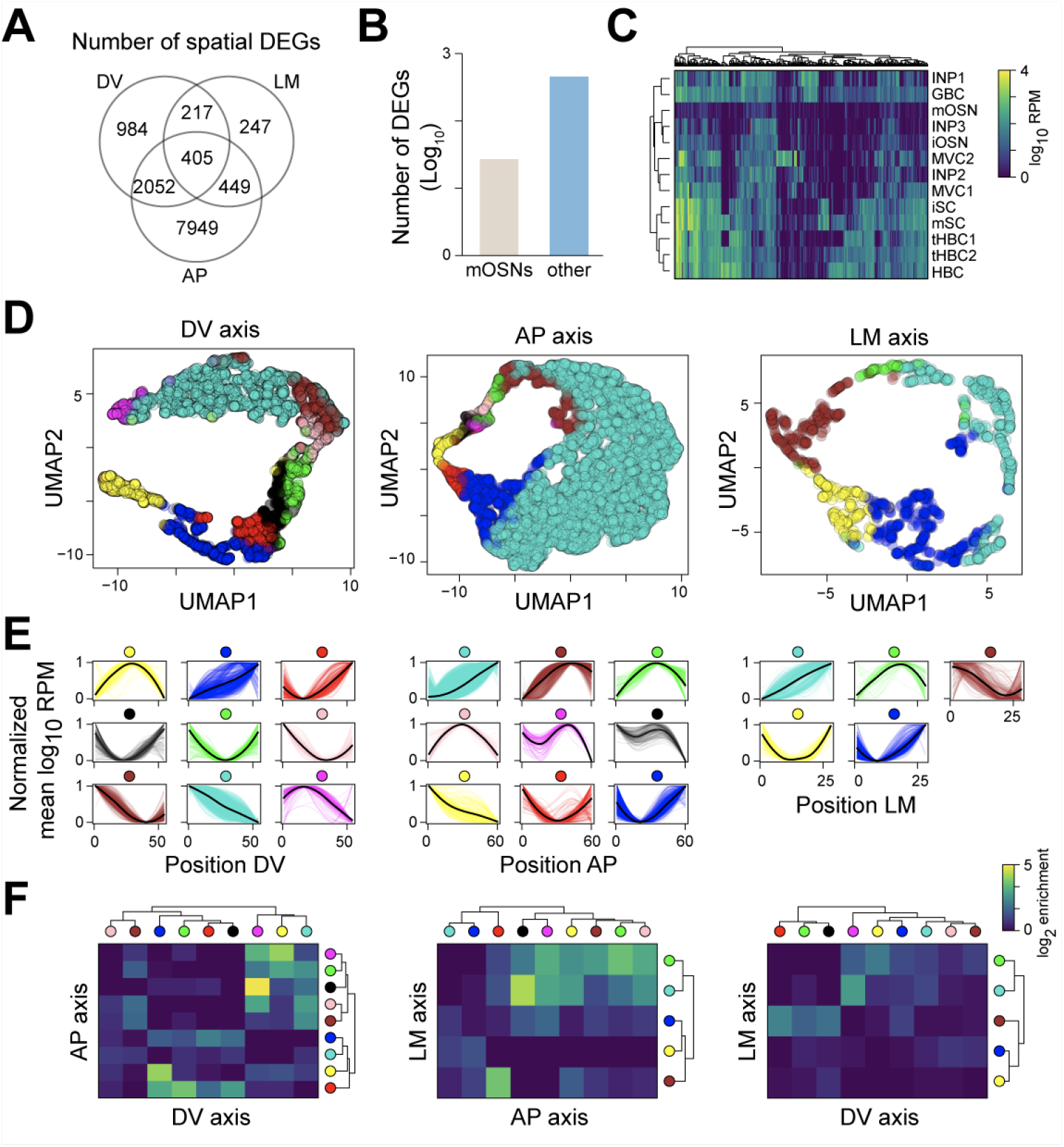
Genes with non-random spatial patterns across different cell types in the OM. (A) Venn diagram showing the numbers of spatial differentially expressed genes (DEGs) along each axis. (B) Bar plot showing the log10 number of spatial DEGs that are mOSN-specific (“mOSNs”), or that are detected only in cell types other than mOSNs (“other”). (C) Heatmap of log10 mean expression per cell type of genes that are not expressed in mOSNs, but only in other OM cell types (INP = Immediate Neuronal Precursors; GBC = Globose Basal Cells; mOSNs = mature Olfactory sensory neurons; iOSNs = immature Olfactory Sensory Neurons; MVC = Microvillous Cells; iSC = Immature Sustentacular Cells; mSC = Mature Sustentacular Cells; HBCs = Horizontal Basal Cells). (D) UMAP plots of spatial DEGs along the three axes. Each gene is colored according to the cluster it belongs to. (E) Normalized average expression patterns of spatial DEGs clusters along the three axes. (F) Heatmap showing the log2 enrichment over the random case for the intersection between lists of genes belonging to different clusters (indicated by colored circles) across pairs of axes.

To add cell-type resolution to the spatial axes, we combined our data with a single-cell RNA-seq (scRNA-seq) dataset from 13 cell types present in the mouse OM (*54*). We catalogued spatial DEGs based on their expression in mature OSNs (mOSNs) versus the 12 other cell types (non-mOSNs) (Figures 2B and 2C; Table S2). This led to the identification of 456 spatial DEGs expressed exclusively in non-mOSNs, which are associated with gene ontology (GO) terms such transcription factors, norepinephrine metabolism, toxin metabolism, bone development, regulation of cell migration, T-cell activation, and others (Table S2). Some of these genes are expressed across many cell types, but others are specific to a single cell type (Figure 2C; Table S2). As expected, among these genes we find some known cell-specific markers with spatial expression patterns, such as the sustentacular cells and Bowman’s glands markers *Cyp2g1* and *Gstm2* (*36*), the neural progenitor cell markers *Eya2* and *Hes6* (*45*), and the basal lamina and olfactory ensheathing cell markers *Aldh1a7* and *Aldh3a1* (*47*) (Table S2). Additionally, we identified many new genes with zonal expression patterns along a single axis or multiple axes, and specific to one or few cell-types (Figures S2B and S2C).

For example, the ribosomal protein *Rps21* plays a key role in ribosome biogenesis, cell growth and death (*55*), and is primarily expressed in horizontal basal cells (HBCs), consistent with the role of HBCs in the maintenance and regeneration of the OE (*56*). Another example is the extracellular proteinase inhibitor *Wfdc18,* which induces the immune system and apoptosis (*57*) and is expressed in microvillous cells type 1 (MVC1s), consistent with the role that MVC1s play in immune responses to viral infection (*58*). Two more examples are the fibroblast growth factor *Fgf20* in immature sustentacular cells (iSCs), and the adapter protein *Dab2* in mature sustentacular cells (mSCs) (Figures S2B and S2C). The fibroblast growth factor Fgf20 is expressed in several cell types, regulates the horizontal growth of the olfactory turbinates, and is preferentially expressed in the lateral OM (*59*), consistent with our data. The adapter protein Dab2 regulates mechanisms of tissue formation, modulates immune responses, and participates in the absorption on proteins (*60, 61*), consistent with the known maintenance and support roles of mSCs in the OM (*62*).

A gene ontology (GO) enrichment analysis on the axis-specific DEGs for non-mOSNs genes revealed a very wide variety of biological processes/molecular functions. Some of the notable terms identified were water and fluid transport (e.g., *Ctfr*, *Aqp3*, *Aqp5*), transcription factors (e.g., *Hes1*, *Hey1*, *Dlx5*), oxidation-reduction processes (e.g., *Scd2*, *Cyp2f2*, *Cyp2g1*), microtubule cytoskeleton organization involved in mitosis (e.g., *Stil*, *Aurkb*), cell cycle (e.g., *Mcm3*, *Mcm4*, *Mcm6*), cell division (e.g., *Kif11*, *Cdca3*, *Nde1*), negative regulation of apoptosis (e.g., *Dab2*, *Scg2*, *Id1*), sensory perception of chemical stimulus, cellular processes (e.g., *Mal*, *Pthlh, Cdc16*), among many others (Table S2).

The successful identification of thousands of spatial DEGs, prompted us to examine their distribution patterns along each axis, and the putative functions associated with such spatial clusters of gene expression. We started by using uniform manifold approximation and projection (UMAP) (*63*) and hierarchical clustering to visualize and cluster all spatial DEGs along the three cartesian axes. This analysis uncovered nine patterns of expression in the DV and AP axes each, and five patterns in the LM axis (Figure 2D-E). These patterns include variations of four major shapes: monotonically increasing (/), monotonically decreasing (\), U-shape (∪), and inverted U-shape (∩) (Figure 2E). The latter two patterns present clear maximum/minimum at different positions along the axis – for example, the brown, green, pink, magenta, and black AP clusters show a similar inverted ∪ pattern, but their maximum moves along the axis (Figure 2E). As expected, several known genes with zonal expression patterns are expressed in the cluster mimicking their respective expression pattern in the tissue. For example, the dorsomedial markers *Acsm4* and *Nqo1* belong to the turquoise clusters in both the DV and LM axes, while the ventrolateral marker *Reg3g* belongs to the blue cluster from the DV axis (Figures 1F and S1F; Table S3).

The total number of genes per cluster had a median value of 236, but varied greatly between clusters – ranging from 57 in the green LM cluster to 8,551 in the turquoise AP cluster (Figure 2D; Table S3. GO enrichment analysis on the axis-specific DEGs yielded enriched terms for 14 of the 23 spatial clusters (Table S3). For example, the turquoise AP cluster displaying a monotonically increasing pattern (Figure 2E) yielded GO terms broadly associated with the molecular machinery of mOSNS – such as axonal transportation, RNA processing, protein modification and quality control, ribosomal regulation and regulation of histone deacetylation (Figure S2D; Table S3). Interestingly, the brown DV cluster, which displays a monotonically decreasing expression pattern (Figure 2E), had similar GO term enrichment (Figure S2E; Table S3). These results raise the hypothesis that mOSN activity is enriched in the dorsoposterior region of the OM, consistent with previous results (*64, 65*).

We then extended our analysis to the remaining clusters, and found additional GO terms shared between several clusters among the three different axes. For example, the GO terms enriched in the dorsomedial region (turquoise DV, pink AP and LM green) suggest that this region is involved in the OM detoxification (Figures S2F-H). Moreover, the enrichment in terms related to immune system in the anteromedial section along the AP axis (yellow, black and magenta AP clusters, and turquoise LM cluster; Table S3) hints at a role of this area in defending OM from pathogenic invaders. Finally, the ventral portion of the DV (red DV cluster) was associated with terms related to cilia movement and function, consistent with both the location and functions of the respiratory epithelium (*36*) (Table S3).

Next, we further explored the relationships between the genes populating each cluster. We found that ventral genes (blue DV cluster) tend to reach a peak in expression in the anterior area of the OM (yellow AP cluster) more often than expected by chance (Figure 2F). We also observed that medial genes (turquoise LM cluster) are more highly expressed in the dorsal (magenta DV cluster) and anterior regions (black, yellow and magenta AP cluster), while genes peaking in the lateral region (brown LM cluster) tend to be ventral (red DV cluster; Figure 2F). These conclusions hold even when we exclude olfactory receptor genes from the analysis (Figures S2I-K).

These associations between the clusters of DEGs along different axes strongly suggest that the presence of complex 3D expression patterns in OM is not restricted to either *Olfrs* or mOSNs. Moreover, our results show that our experimental approach can uncover spatially restricted functional hotspots within the OM.

### A 3D transcriptomic atlas of the mouse OM

Since the OR discovery three decades ago (*2*), in-situ hybridization (ISH) has been the method of choice to study spatial gene expression patterns across the OM. This method can be technically challenging and is inherently a very low-throughput experimental approach.

As we showed above, our Tomo-seq data enables a systematic and quantitative estimation of gene expression levels along the three cartesian axes of the OM. Here, we take this analysis one step further and generate a fully browsable tridimensional (3D) gene expression atlas of the mouse OM. To this aim, first we reconstructed the 3D shape of OM based on publicly available images of OM sections (made available by (*66*), see STAR Methods). We then fed the resulting shape information combined with the gene expression data along the three cartesian axes into the iterative proportional fitting (IPF) algorithm (*30, 67*) (Figure 3A). Our resulting 3D atlas of the OM faithfully reproduced the known 3D pattern of the dorsomedial marker *Acsm4* (*34*) (Figure 3B). To further validate that our 3D atlas recapitulates known patterns of gene expression in the OM, we compared the 3D reconstructed patterns with conventional ISH patterns for one novel spatial DEGs identified in this study, *Cytl1* (Figure 3C and 3D; Table S3). *Cytl1* is expressed mainly along the septum (Figure 3C and 3D), consistent with the role *Cytl1* plays in osteogenesis, chondrogenesis, and bone/cartilage homeostasis (*68, 69*).

**Figure 3.**
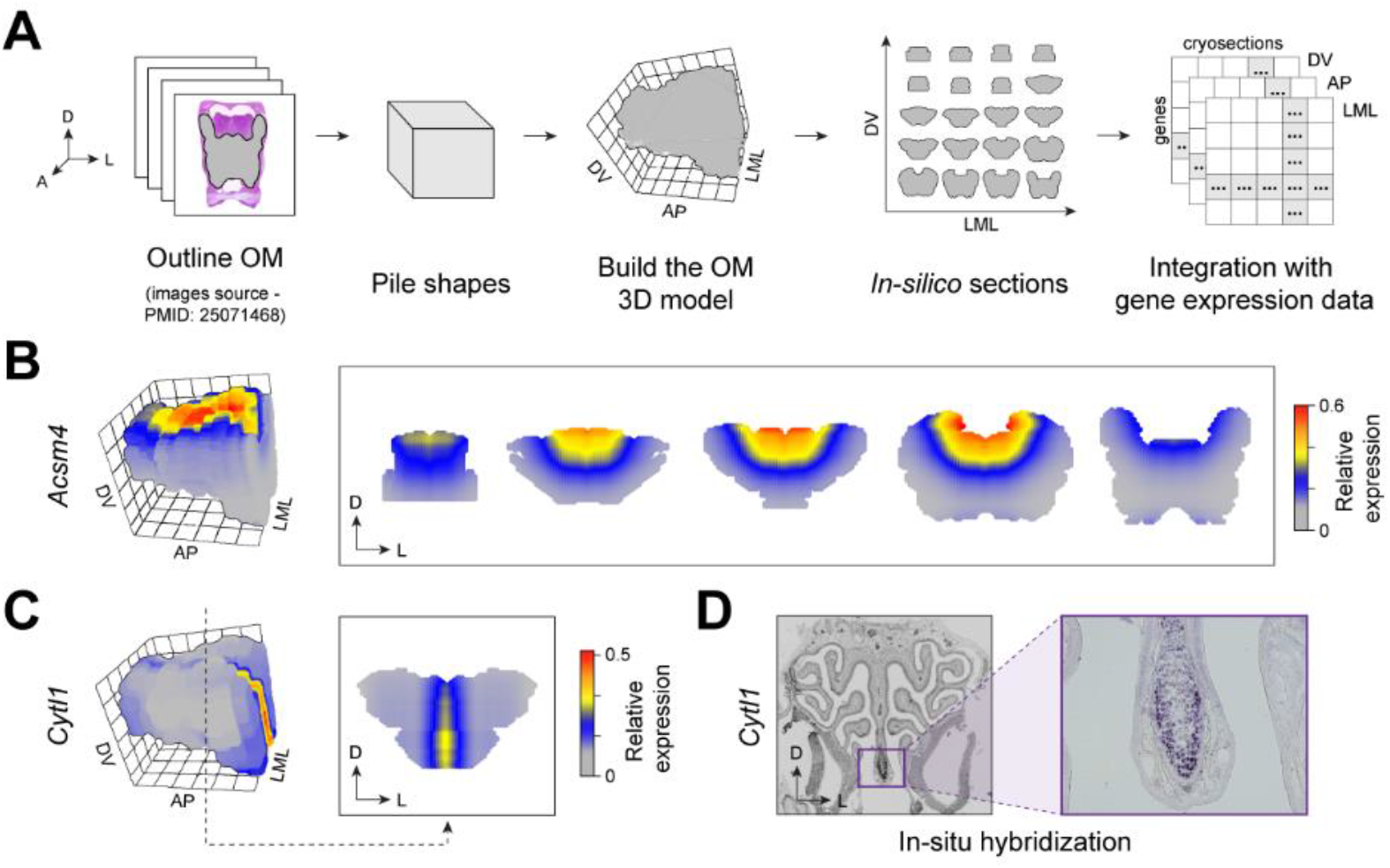
The 3D reconstruction of the OM. (A) Schematic of 3D shape reconstruction strategy. Images of 2D slices along the AP axis of the OM were piled together to build an in-silico 3D model of OM, which can also be used to visualize in silico sections. This 3D model, together with the gene expression data along each axis, was the input of the iterative proportional fitting algorithm, which allowed us to estimate a 3D expression pattern for any gene. (B) Reconstruction of the 3D expression pattern of the gene *Acsm4* in the OM, visualized in 3D and in OM coronal sections taken along the AP axis. (C) Reconstruction of the 3D expression pattern of the gene *Cytl1* in the OM. (D) In-situ hybridization experiment validating *Cytl1* spatial expression pattern reconstructed in panel D.

To make this 3D gene expression atlas of the mouse OM available to the scientific community, we created a web-portal (available at http://atlas3dnose.helmholtz-muenchen.de:3838/atlas3Dnose) providing access to the spatial transcriptomic data described here in a browsable and user-friendly format. Specifically, this portal contains search functionalities that allows the users to perform pattern search by gene, which returns: i) the normalized counts along each of the three cartesian axes; ii) the predicted expression pattern in the 3D OM with a zoom function; iii) visualization of the expression patterns in virtual cryosections along the OM, by selecting any possible pairwise intersection between two given axes (i.e., DV x AP, DV x LM, AP x LM); iv) the degrees of belonging for each ‘zone’ (see results section below); and v) single-cell expression data across 14 different OM cell types (see results section below).

In sum, here we generated and made publicly available a highly sensitive 3D gene expression atlas of the mouse OM that allows the exploration of expression patterns for nearly 20,000 genes.

### Topographical expression patterns of *Olfrs*

Early ISH studies postulated that OSNs expressing different ORs are spatially segregated into four broad areas within the MOE, called ‘zones’, and which define hemicylindrical rings with different radii (*12, 13*). Subsequent studies identified ORs expressed across multiple zones, making clear that a division in four discrete zones might not accurately reflect the system, and a continuous numerical index representing the pattern of expression of each OR along the five OE zones was implemented (*14, 15*). More recently, a study reconstructed OR gene expression patterns in three dimensions (3D), and qualitatively classified the expression areas of 68 OR genes in nine overlapping zones (*16*). However, all these studies combined sampled but a fraction (~10%) of the total intact OR gene repertoire and, most importantly, lack a quantitative and unbiased definition of zones or indices.

In our combined dataset we detected a total of 959 unique *Olfrs* (Figure 4A), of which we confidently reconstructed the spatial expression patterns for 689 differentially expressed in space (FDR<0.01; Figure 4B) – a number six times larger than the combined 112 *Olfrs* characterized by previous ISH studies (*10, 12, 14, 16*). To define *Olfr* expression in 3D space in a rigorous, unbiased and quantitative way, we ran a Latent Dirichlet Allocation algorithm (see STAR Methods)(*70*) on the 689 spatially differentially expressed *Olfr*, which suggested the presence of five zones (Figure S3A; STAR Methods). Next, we visualized the spatial distribution of these five zones in our 3D model of the OM, with colors representing the probability that a given spatial position belongs to each zone (Figure 4C). These zones extend from the dorsomedial-posterior to the lateroventral-anterior region, consistent with the previously described zones (*12–14*).

**Figure 4.**
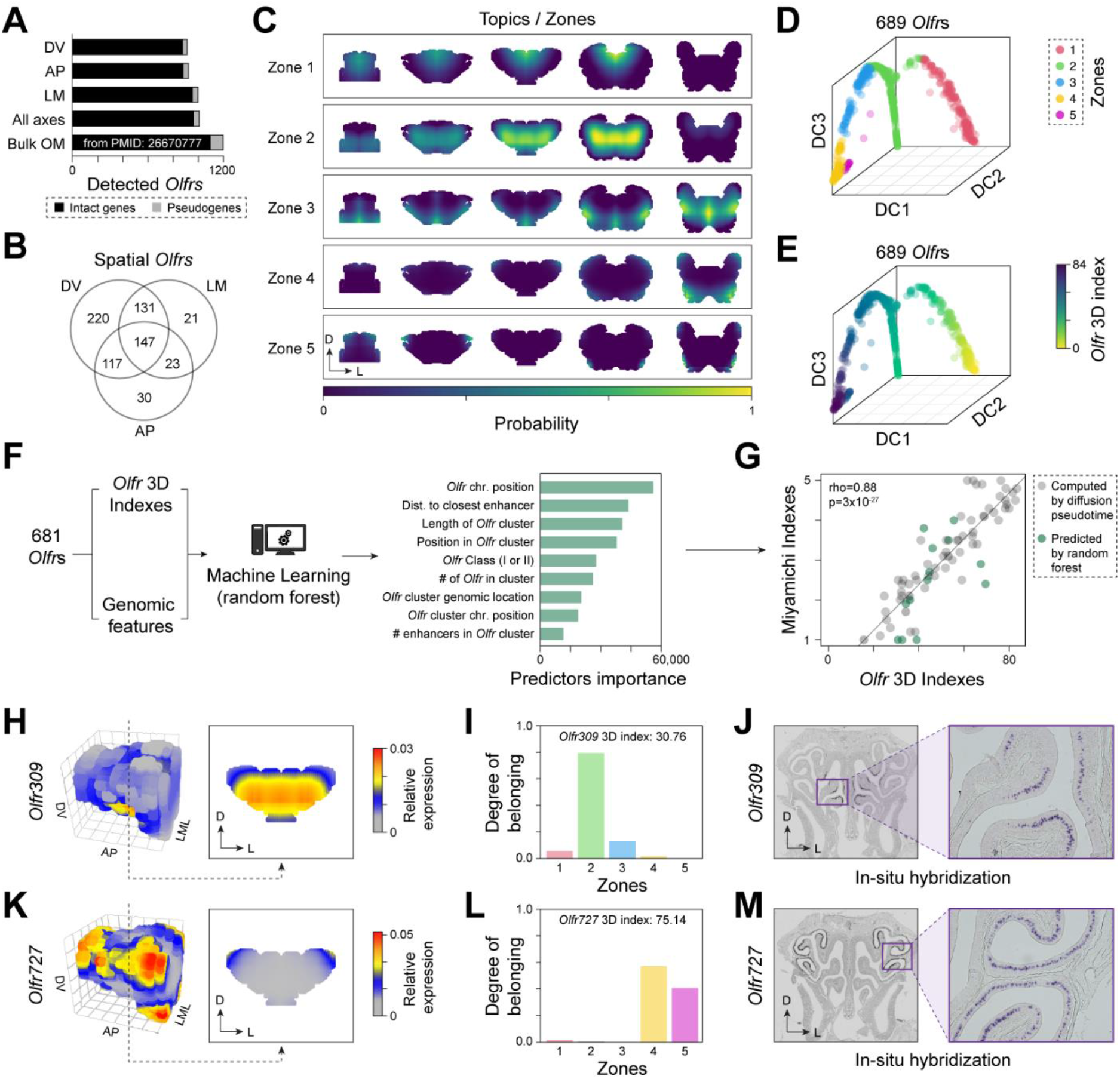
Zonal organization of the OM. (A) Number of Olfr genes detected in our data and in an OM bulk RNA-seq data (6) (B) Venn diagram of spatially differentially expressed Olfrs per axis. (C) Visualization of the five zones across the OM (coronal sections) estimated with a Latent Dirichlet Allocation algorithm. The colors indicate the probability (scaled by its maximum value) that a position belongs to a given zone. (D) Diffusion map of *Olfrs*. Genes are colored based on the zone they fit in the most. DC, diffusion component. (E) Same as panel D, with Olfrs colored by their 3D index. (F) We fitted a Random Forest algorithm to the 3D indexes of 681 spatially differentially expressed *Olfr* using nine genomic features as predictors. After training, the Random Forest was used to predict the 3D indexes of 697 *Olfrs* that have too low levels in our data. (G) 3D indexes versus the indexes of 80 *Olfrs* estimated in (14) from ISH data. Black circles indicate *Olfrs* detected in our dataset; green circles are *Olfrs* whose indexes were predicted with Random Forest. (H, K) Predicted expression patterns of *Olfr309* and *Olfr727*. (I,L) Degrees of belonging for *Olfr309* and *Olfr727*. (J,M) In-situ hybridization for *Olfr309* and *Olfr727*.

The majority of *Olfrs* with known spatial patterns are restricted to one zone, but a small number of *Olfrs* are expressed across multiple zones, in a continuous or non-continuous fashion (*14–16*). Under this logic, each *Olfr* has a different probability of belonging to the five different topics/zones we identified. To test this assumption, we used the same mathematical framework as above to compute the probabilities that the expression pattern of each *Olfr* gene belongs to a given zone, i.e., the “degree of belonging” (DOB, Table S4). The DOBs represent a decomposition of the expression patterns in terms of the five zones (Figure 4C) and quantitatively describe the changes in patterns of genes with overlapping areas of expression (e.g., see Figure S3B). The width of the distribution of DOBs across the five zones, which can be measured with entropy, can distinguish genes whose patterns mostly fit in a single zone from those spanning multiple zones (Figure S3C; STAR Methods).

To visualize the global distribution of the 689 *Olfrs*, we applied the diffusion map algorithm (*71*) to their DOBs. This showed that the genes are approximately distributed along a continuous line spanning the five zones and without clear borders between zones (Figure 4D) (*14–16*). With the diffusion pseudo-time algorithm (*72*), we calculated an index (hereafter referred to as “3D index”) that tracks the position of each *Olfr* gene along the 1D curve in the diffusion map and represents its expression pattern (Figure 4E).

While our approach yielded an index for the 689 spatially differentially expressed *Olfr* genes used to build the diffusion map, there were ~700 *Olfrs* that could not be analyzed, either because they were too lowly expressed or not detected at all in our dataset (Figure 4A). However, since the spatial expression patterns for some *Olfrs* are partly associated with their chromosomal/genomic coordinates (*73–75*), we hypothesized that we could train a machine learning algorithm to predict the 3D indices for the ~700 *Olfrs* missing from our dataset. Thus, we trained a Random Forest algorithm on the 3D indices of the spatially differentially expressed *Olfrs* in our dataset using 9 genomic features as predictors (e.g., chromosomal position, number of *Olfrs* in cluster, distance to nearest known enhancer, etc; see STAR Methods and Figure 4F). The five most important predictors were features associated with chromosomal position, distance to the closest *Olfr* enhancer (*76*), length of the *Olfr* cluster, position in the *Olfr* cluster, and phylogenetic class of *Olfrs* (Figure 4F).

After assessing the performance of the algorithm with a cross-validation test (Figure S4A; STAR Methods), we predicted the 3D indices for the 697 *Olfrs* missing reliable expression estimates in our dataset. Overall, through multiple unsupervised and supervised computational methods, we have quantitatively defined five spatial expression domains in the OM (called zones), and have provided accurate 3D spatial indices for 1386 *Olfrs*, which represents ~98% of the annotated *Olfrs*.

Importantly, we found that our 3D indices strongly correlate with the zonal indices inferred using ISH in (*14*)(rho=0.88, p=3×10^−27^; Figure 4G) and in two other studies (*16, 73*)(Figures S4B and S4C). Furthermore, we performed ISH for two *Olfrs* that have not been characterized before, *Olfr309* and *Olfr727*, which were correctly predicted to be expressed primarily in zone 2 and 4 respectively (Figures 4H-M; Table S4).

### Topographical expression patterns for non-Olfr genes

Using the mathematical framework based on topic modelling described above, we also decomposed the expression patterns of non-*Olfr* genes onto the five zones we identified. This gave us the opportunity to look for genes showing zone specificity by calculating the entropy of the DOBs distributions (see above). Interestingly, we found 28 genes that are highly specific for each of the five zones (i.e., with entropy <1; see Figure 5A, STAR Methods and Table S5). For example, *S100a8* (zone 1) codes for a S100 calcium-binding protein A8 protein involved in calcium signaling and inflammation (*77*); *Moxd2* (zone 2) is a monooxygenase dopamine hydroxylase-like protein possibly involved in olfaction (*78*); *Lcn4* (zone 3) is a lipocalin involved in transporting odorants and pheromones in the mouse nose (*79, 80*); *Gucy1b2* (zone 4) is a soluble guanylyl cyclase oxygen and nitric oxide (*81, 82*); and *Odam* (zone 5), a secretory calcium-binding phosphoprotein family member involved in cellular differentiation and matrix protein production, and with antimicrobial functions of the junctional epithelium (*83, 84*) (Figure 5B). The high zone-specificity of the expression pattern of these genes gives clues into possible biological processes taking place in the zones. Indeed, *Gucy1b2* is a known genetic marker for a small OSN subpopulation localized in cul-de-sac regions in the lateral OM, consistent with our reconstruction (Figure 5B), and it regulates the sensing of environmental oxygen levels through the nose (*6, 85*). Additionally, we performed in situ hybridization experiments on *Moxd2*, which revealed that it is expressed mostly in a small ventrolateral patch of the OM (Figure 5C and 5D), validating its predicted 3D spatial pattern (Figure 5B), and highlighting a highly localized putative role of this monooxygenase dopamine hydroxylase-like protein in neurotransmitter metabolism (*86*) in the mouse OM.

**Figure 5.**
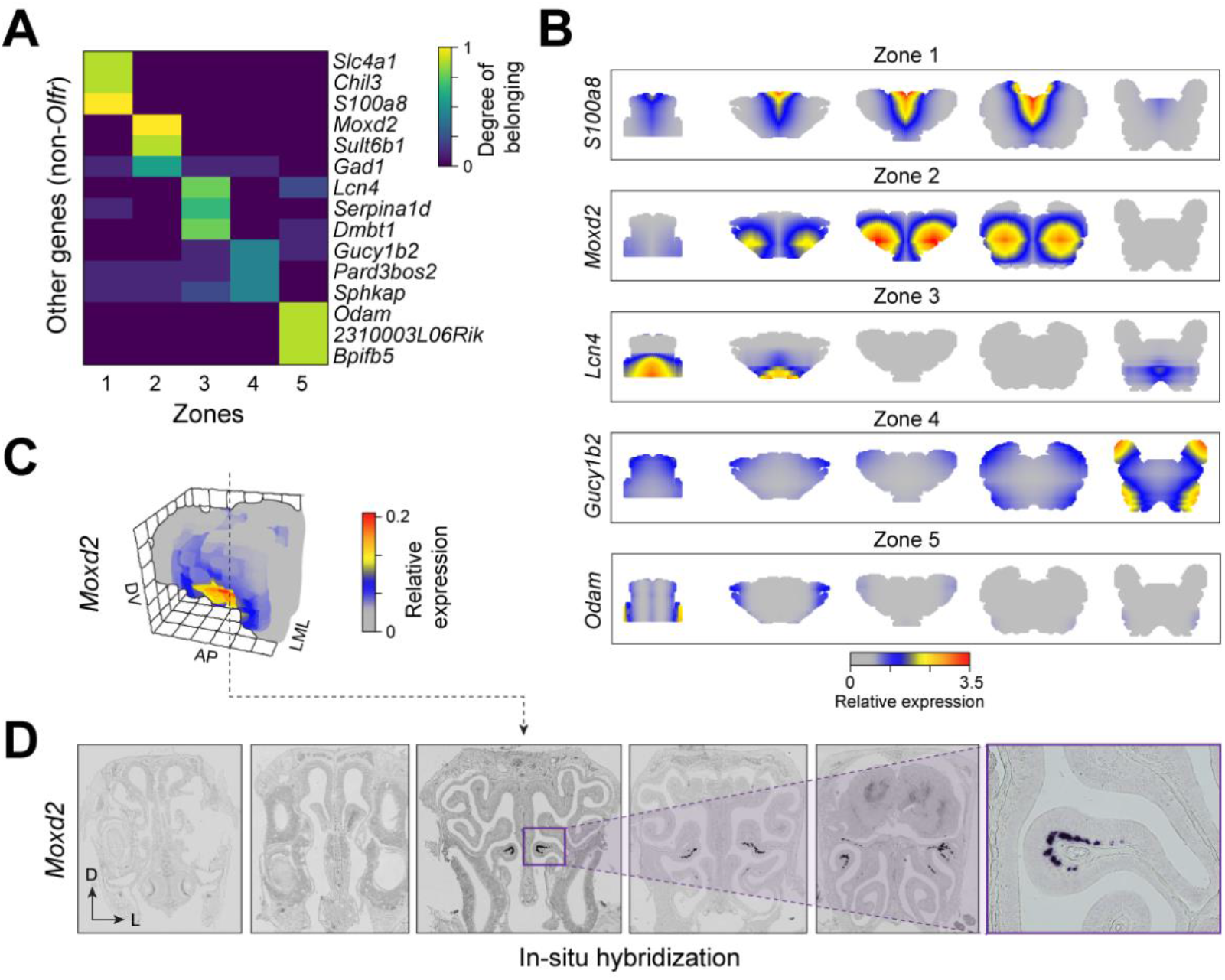
Non-Olfr genes. (A) Heatmap of degrees of belonging of most zone-specific non-*Olfr* genes. (B) 3D gene expression pattern (coronal sections) of most topic specific non-*Olfrs* for each topic along the AP axis. (C) Reconstruction of the 3D expression pattern of the gene *Moxd2* in the OM. (D) In-situ hybridization experiment validating *Moxd2* spatial expression pattern reconstructed in panel C.

A recent study showed that the transcription factors Nfia, Nfib and Nfix regulate the zonal expression of *Olfrs* (*17*). To get some insights into the signaling pathways involved in this process, we mined our dataset for genes encoding ligands and receptors (*87*) that significantly correlate with the expression patterns of the Nfi transcription factors (see STAR Methods). This analysis returned 476 genes involved in biological processes primarily associated with regulation of neurogenesis, regulation of cell development, regulation of nervous system development, anatomical structure development, cellular component organization or biogenesis and regulation of neuron differentiation (Table S5). As expected, some of these genes have known functions in the OM, such as segregating different cell lineages in the OM for *Notch1-3* (*88*), genes associated with the development of the nervous system (e.g., *Erbb2* and *Lrp2*) (*89, 90*), and many others associated with the semaphorin-plexin, ephrin-Eph, and Slit-Robo signaling complexes – which regulate OSN axon guidance and spatial patterning of the OM (*42, 91–93*). Excitingly, the majority of these 476 genes still have unknown functions in the OM, thus highlighting the potential of our approach to discover new genes and pathways involved in the regulation of zonal expression in the OM.

### The anatomical logic of smell

For most sensory systems, the functional logic underlying the topographic organization of primary receptor neurons and their receptive fields is well-known (*1*). In contrast, the anatomic logic of smell still remains unknown, and it is subject of great controversy and debate (*23, 26*).

To explore the underlying logic linked to Olfrs zonal distribution, we investigated possible biases between the expression patterns of Olfrs and the physicochemical properties of their cognate ligands. First, we compiled a list of known 738 Olfr-ligand pairs, representing 153 Olfrs and 221 odorants (Figure 6A; Table S6). Interestingly, we found that *Olfr* pairs sharing at least one common ligand have more similar expression patterns (i.e., more similar 3D indices) than Olfrs detecting different sets of odorants (Figure 6B). This observation is consistent with the hypothesis that the *Olfr* zonal expression depends, at least partially, on the properties of the odorants they bind to.

**Figure 6.**
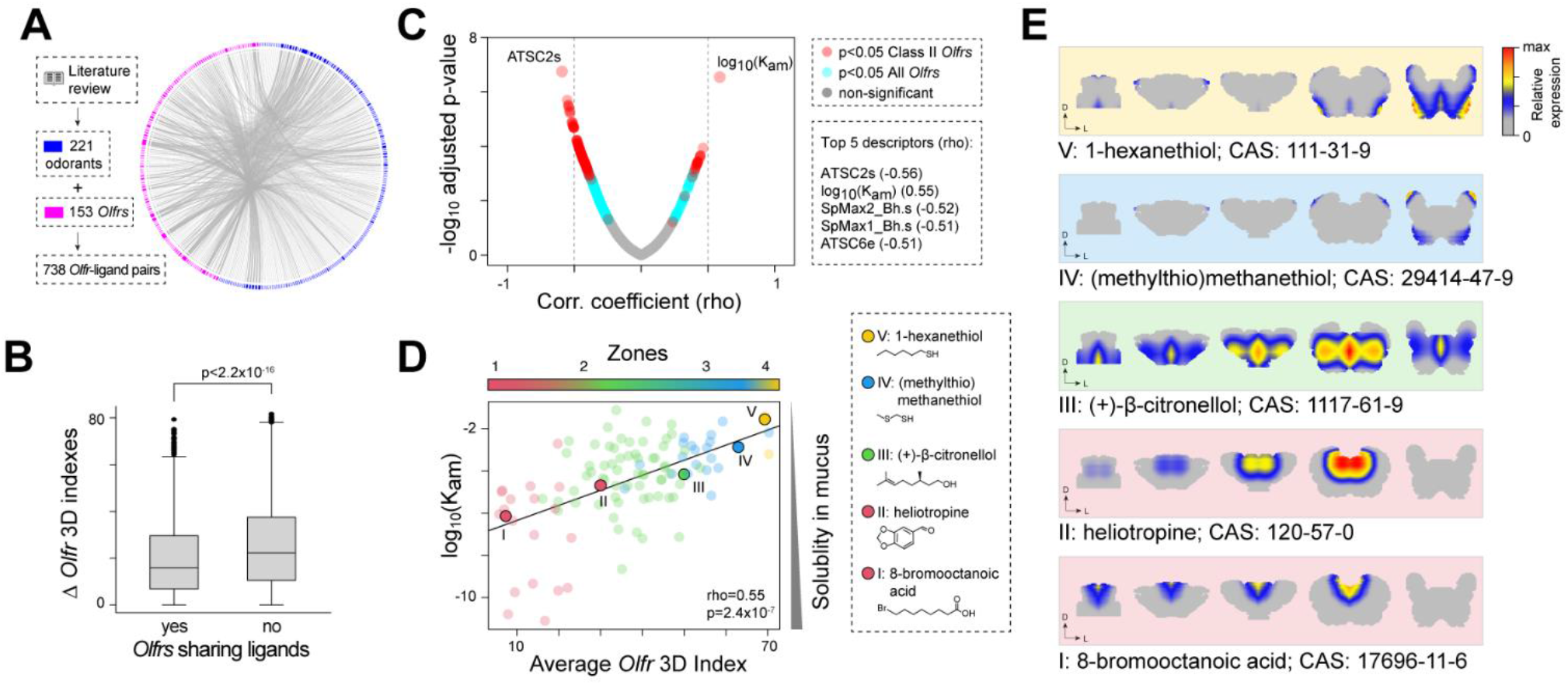
Physiological role of the zones. (A) Circular network illustrating the pairs of Olfrs and ligands that we found in literature. (B) Box plots showing the distributions of the absolute value of 3D index differences between pairs of Olfrs sharing at least one ligand versus pairs of Olfrs without cognate ligands in common. The difference between the two distributions is statistically significant (p < 2.2×10^−16^, Wilcoxon Rank-Sum test) (C) Scatter plot showing the Spearman correlation coefficients between the ligands’ mean 3D indices and molecular descriptors, and the corresponding −log_10_(adjusted p-value). Turquoise circles indicate the descriptors having a significant correlation only when both class I and II Olfr are considered; red circles mark the descriptors with a significant correlation also when class I Olfr are removed. (D) Scatter plot illustrating the correlation between air/mucus partition coefficients of the odorants and the average 3D indexes of their cognate Olfrs. Only odorants for which we know at least two cognate Olfrs (110) were used here. Odorants are colored according to the zone they belong to (defined as the zone with the highest average degree of belonging computed over all cognate receptors). The five odorants highlighted in the plot by larger circles are indicated on the right-hand side, along with their molecular structure and common name. (E) Average expression pattern of the cognate Olfrs recognizing each of the five odorants highlighted in panel D, including their respective CAS numbers.

Next, we considered a set of 1210 physicochemical descriptors, including the molecular weight, the number of atoms, aromaticity index, lipophilicity, and the air/mucus partition coefficient (*K_am_*), which quantifies the mucus solubility of each ligand (*21, 94*) (see STAR Methods). We then computed the Spearman’s correlation of each of these descriptors of the ligands with the average 3D indices of the Olfrs detecting them (see STAR Methods). We found a statistically significant correlation for 744 descriptors (FDR < 0.05 see Figure 6C). The highest correlation is with the air/mucus partition coefficient *K_am_* (rho = 0.55, adjusted p-value = 2×10^−7^). Interestingly, among the top five correlating descriptors, three more (ATSC2S, SPmax1_Bh.s and SPmax2_Bh.s; Figure 6C) are also related to solubility (*95–97*). These correlations are robust to changes in the set of ligands and/or Olfrs used for the analysis (e.g., when restricting the analysis only to class II Olfrs, see STAR Methods).

In particular, the positive correlation of the 3D indices with *K_am_* (see Figure 6D) indicates that the most soluble odorants (lower *K_am_*) preferentially activate Olfrs expressed in the most antero-dorsomedial OM region (zone 1) of the OM, while the least soluble odorants (higher *K_am_*) activate Olfrs in the postero-ventrolateral OM region (zones 4-5). In other words, gradients of odorants sorption (as defined by their *K_am_*) correlate with the gradients of Olfr expression from zone 1 to zone 5, consistent with the *chromatographic/sorption* hypothesis in olfaction (*20, 21*).

This is exemplified by the plots in Figure 5D, illustrating the predicted average expression levels across OM sections of the Olfrs binding to five odorants with different values of *K_am_*. These results show, for the first time, a direct association between *Olfr* spatial patterns and the calculated sorption patterns of their cognate ligands in the OM, providing a potential explanation for the physiological function of the zones in the OM.

## DISCUSSION

Here we presented a 3D high-resolution transcriptomic atlas of the mouse OM, which allows, for the first time, the exploration and visualization of the expression patterns for thousands of genes. By integrating our 3D atlas with a previously published single-cell RNA-sequencing dataset (54), we were also able to quantify the expression of these genes across 14 different cell types populating the mouse olfactory mucosa. To facilitate the exploration and dissemination of this important resource, we developed a powerful yet easy to use online browser that allows gene queries returning 3D and single-cell specific gene expression patterns in a user-friendly graphical interface. This atlas allowed us to identify spatial blueprints and provided further insight into the functional logic underlying the molecular organization of the mouse olfactory peripheral system.

Past studies yielded inconclusive and sometimes contradictory views on the basic logic underlying the peripheral representation of smell, partly because the topographic distribution of OSN subtypes and their receptive fields still remained vastly uncharted, data on Olfr-ligand pairs was scarce, and the many pitfalls associated with electro-olfactogram recordings used to study spatial patterns of odor recognition in the nose (26, 98, 99). Here, we combined RNA-seq and computational approaches that utilize unsupervised and supervised machine learning methods to discover and quantitatively characterize spatial expression patterns in the OM. We created the first 3D transcriptional map of the mouse OM, which allowed us to spatially characterize 17,628 genes, including ~98% of the annotated Olfrs. We identified and validated by ISH several new spatial marker genes, and a clustering analysis pinpointed the OM locations where specific functions related to, e.g., the immune response, might be carried out. We also identified five broad Olfr expression zones in the OM, which were mathematically defined and used to decompose the expression patterns of all genes.

Our analysis also enabled us to answer fundamental and longstanding questions about the rationale behind the spatial organization of the peripheral olfactory system. Specifically, we provide evidence to the hypothesis that the spatial zones increase the discriminatory power of the olfactory system by distributing Olfr receptors in the areas of the OM that are more likely to be reached by their cognate ligands, based on their solubility in mucus. However, a caveat of this approach is that the Olfr-ligand list we compiled from the literature includes odorant libraries of different size and composition, and tested using different experimental approaches. Moreover, highly abundant Olfrs have a higher probability of being deorphanized than lowly abundant Olfrs, and ecologically relevant odorants are more likely to activate Olfrs when compared to other odorants (100–102). Despite having compiled and performed our analysis on the largest set of Olfr-ligand pairs assembled to date and carrying out multiple robustness checks, we cannot rule out that ascertainment bias might contribute to the associations we found between the Olfr spatial location and the properties of their respective ligands. Future studies investigating the activation profiles for all mouse Olfrs and/or mapping the in-vivo activation patterns of mouse Olfrs in the olfactory mucosa will be key to stress test the conclusions of our study.

The quantitative framework we built for this dataset will facilitate interrogation of gene expression patterns via an online tool we provide, and help answer important questions on the spatial patterns in the nose. Moreover, our approach could be easily applied to spatial transcriptomic data collected from other tissues, or the same tissue across multiple developmental stages. Results from this study will also serve as a template to start answering other important questions about olfaction, such as whether Olfr spatial expression maps can also encode maps of odor perception. Because the general molecular mechanisms of olfaction, zonal organization of Olfrs, and components of olfactory perception are conserved in mammals (100, 103-106), findings from our and other subsequent studies can likely be extrapolated to other mammals, including humans. Finally, the functional logic underlying the topographic organization of primary receptor neurons and their receptive fields in smell is now starting to be exposed.

## ACKNOWLEDGMENTS

We would like to thank Maria-Elena Torres-Padilla, Bob Datta, and the members of the Saraiva and Scialdone Labs for the constructive feedback, and helpful comments. We are thankful to Giorgia Greco for insightful discussions about chemical properties of ligands. This work was supported by the Helmholtz Association (Scialdone Lab) and Sidra Medicine (project #SDR400040 to L.R.S.).

## AUTHOR CONTRIBUTIONS

M.L.R.T.S. analyzed data and wrote the initial version of the manuscript. E.A.M. performed the RNA-seq experiments and analyzed data. T.S.N., L.S.M., and S.L. performed experiments. M.M., S.S.Y.H., J.D.M., F.V., M.C., and M.O. analyzed data. E.G., J.R., D.W.L., and B.M. analyzed data, and helped write the manuscript. A.S. and L.R.S. conceived and supervised the project, analyzed data, and wrote the final version of the manuscript.

## MATERIAL AND METHODS

### Animals

The animals used in this study were adult male C57Bl/6J mice (aged 8-14 weeks, The Jackson Laboratory, Stock # 00664) maintained in group-housed conditions on a 12:12 h light:dark schedule (lights on at 0700 hours). Each mouse was randomly assigned for cryosectioning along one of the three cartesian axes.

The use and care of animals used in this study was approved by the Internal Animal Care and Use Committee (IACUC) of Monell Chemical Senses Center, by the IACUC of the University of São Paulo, and by the Wellcome Trust Sanger Institute Animal Welfare and Ethics Review Board in accordance with UK Home Office regulations, the UK Animals (Scientific Procedures) Act of 1986.

### Dissection of the olfactory mucosa, cryosections, and RNA-sequencing

The olfactory mucosa (OM) of 9 mice was carefully dissected, and all the surrounding tissue (including glands and bone) removed. The OMs were then embedded in OCT (Tissue Tek), immediately frozen in dry-ice and kept at −80°C. Each OM was then cryosectioned along each of the 3 cartesian axes: dorsal-ventral (DV, N=3), anterior-posterior (AP, N=3), or lateral-medial-lateral (LML, N=3). Every second cryosections (35 ***μ***m thick) was collected into 1.5 mL eppendorf tubes containing 350 ***μ***l RLT Plus Buffer (Qiagen) supplemented with 1% 2-mercaptoethanol, immediately frozen in dry-ice and kept at −80C until extraction. RNA was extracted using the RNeasy Plus Micro Kit (Qiagen), together with a genomic DNA eliminator column and a 30-minute incubation with DNAse I (Qiagen). Reverse transcription and cDNA pre-amplification were performed using the SMART-Seq v4 Ultra Low Input RNA Kit for Sequencing (Clontech/Takara). cDNA was harvested and quantified with the Bioanalyzer DNA High-Sensitivity kit (Agilent Technologies). Libraries were prepared using the Nextera XT DNA Sample Preparation Kit and the Nextera Index Kit (Illumina). Multiplexed libraries were pooled and paired-end 150-bp sequencing was performed on the Illumina HiSeq 4000 platform at Sidra Medicine, except for one library (DV-I) for which 125-bp paired-end sequencing was performed on the Illumina HiSeq 2500 platform at the Wellcome Sanger Institute. For the remaining eight libraries, 150-bp sequencing was performed on the Illumina HiSeq 4000 platform. The raw data are available through ArrayExpress under accession number E-MTAB-10211.

### RNA-seq data mapping and gene counting

Reads were aligned to the mm10 mouse genome (release 99). The sequences of the genes “*Xntrpc*” and “*Capn5*” were removed from the genome files as in (*6*). The alignment was performed with the software STAR version 2.7.3a (*107*). Genome indexes were generated using STAR --runMode genomeGenerate with default parameters. Then, alignment of reads was performed with the following options: --runThreadN 48 --outSAMunmapped Within --outFilterMultimapNmax 1000 --outFilterMismatchNmax 4 --outFilterMatchNmin 100 --alignIntronMax 50000 --alignMatesGapMax 50500 --outSAMstrandField intronMotif --outFilterType BySJout. The resulting SAM files were converted to bam format and sorted using samtools (version 0.1.19-44428cd)(*108*). The multi mapping reads were eliminated using the same software (samtools view -q 255). Finally, the reads for each gene were counted using htseq-count (version 0.11.2) with the options -m intersection-nonempty -s no -i gene_name -r pos (*109*).

### Quality Control

We excluded all the samples that fulfilled any of these criteria: they had less than 50% mapped reads, less than 4,000 detected genes, more than 20% mitochondrial reads, less than 10,000 total number of reads, or did not express any of the 3 canonical OSN markers *Omp*, *Cnga* and *Gnal*. This resulted in ~51 good-quality sections along the DV axis (~84% out of the collected sections), ~76 (~91% of total) along the AP axis and ~59 (~93% of total) along the LML axis, as averaged across the three replicates per axis.

### Data normalization

Gene expression counts were normalized by reads-per-million (RPM), then genes detected in only one replicate and genes that were detected in less than 10% of all samples along one axis were eliminated. To check the similarity between replicates, we calculated Spearman correlations between the transcriptional profiles of sections along each axis (using the top 1000 Highly Variable Genes per axis). Close positions had the most similar transcriptional profiles (Figure 1C). Afterwards, the 3 replicates for each axis were aligned as follows: the top 3,000 highly variable genes (HVGs) from each replicate were identified using the method implemented in the scran library in R (*110*) and the intersection of these 3 groups was used in the next steps. For the replicates’ alignment, we took as reference the replicate with the smallest number of slices. We used a sliding window approach that identified the range of consecutive positions on each replicate along which the average value of the Spearman’s correlation coefficient computed with the reference replicate over the HVG was maximum (mean Spearman’s Rho= 0.80, p<0.05). To mitigate batch effects, the level of every gene was scaled in such a way that their average value in each replicate was equal to the average calculated across all replicates. After this scaling transformation, the data was then averaged between replicates. Once the 3 biological replicates were combined, we had 54 sections along the DV axis, 60 along the AP and 56 along the LML. Along the LML axis a symmetric pattern of expression is expected around the central position, where the septal bone is located. To confirm this in our data, first we identified the central position by analyzing the expression pattern of neuronal markers like *Cnga2*, *Omp* and *Gnal*, whose expression is lowest in the area around the septal bone. Indeed, all three marker genes reach a minimum at the same position along the LML axis (slice 28), which we considered to be the center. The expression patterns of ~90% of genes on either side of the central position show a positive correlation, and ~70% reach statistical significance (Spearman’s correlation computed on the highly variable genes having more than 50 normalized counts in at least 3 slices), further supporting the hypothesis of the bilateral symmetry. Hence, after replicates were averaged, LML axis was made symmetric averaging positions 1:28 and 56:29. Moreover, *Olfrs* were normalized by the geometric mean of neuronal markers *Omp*, *Gnal* and *Cnga2*, as done previously (*11*).

To verify the presence of a spatial signal, we calculated the Moran’s I and the associated p-values for the top 100 Highly Variable genes along each axis using the “Moran.I” function from the “ape” library in R with default parameters (*111*). The p-values of the genes along each axis were combined with the Simes’ method (*112*) using the function combinePValues from the scran R library (Figure S1E).

### Identification of differentially expressed genes and gene clustering

Before testing for differential expression along a given axis, we filtered out genes whose expression levels had low variability. To this aim, for each gene we estimated their highest and lowest expression by taking the average of its three highest and three lowest values respectively. Then, we considered for downstream analyses only the genes that meet either of these two criteria: the highest expression value is greater than or equal to 5 normalized counts and the fold-change between the highest and lowest value is greater than 2; or the difference between the highest and the lowest value is greater than or equal to 4 normalized counts.

The expression levels of the genes were binarized according to whether their value was higher or lower than their median expression along the axis. Finally, we used the “ts” function in R to transform the binarized expression values into time series objects, and we applied on them the Ljung-Box test (Box.test function in R with lag=(axis length)-10) which identifies genes with statistically significant autocorrelations, i.e., with non-random expression patterns along an axis. The resulting p-values were adjusted using the FDR method and genes with an FDR < 0.01 were considered as differentially expressed. For the next steps, the log10 normalized expression of differentially expressed genes along each axis was fitted with a local regression using the locfit function in the R library locfit (*113*). Smoothing was defined in the local polynomial model term of the locfit model using the function “lp” from the same library with the following parameters: nn = 1 (Nearest neighbor component of the smoothing parameter) and deg = 2 (degree of polynomial). The fitted expression values of these genes along each axis were normalized between 0 and 1. Clustering was performed separately for each axis on the fitted and normalized patterns of the differentially expressed genes. We used the R function “hclust” to perform hierarchical clustering on the gene expression patterns, with a Spearman’s correlation-based distance (defined as 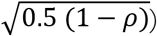) and the “average” aggregation method. The number of clusters were defined with the cutreeDynamic function from the dynamicTreeCut R library, with the parameters minClusterSize = 50, method = “hybrid” and deepSplit = 0. To visualize the data in two dimensions, we applied the UMAP dimensionality reduction algorithm (umap function in the R library umap with default options; see Figure 2D) (*63, 114*). To analyze the relationship between the expression patterns of genes along different axes, we computed the intersections of the gene clusters between any pair of axes. The expected number of elements in each intersection under the assumption of independent sets is given by:

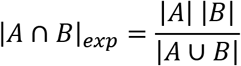

where A and B indicate the sets of genes in two clusters identified along two different axes and |⋅| indicates the cardinality of a set (i.e., the number of its elements). The ratio between the observed and the expected number of elements in the intersection |*A* ∩ *B*|_*obs*_ / |*A* ∩ *B*|_*exp*_ quantifies the enrichment/depletion of genes having a given pair of patterns across two axes with respect to the random case. The log2 values of (1 + |*A* ∩ *B*|_*obs*_ / |*A* ∩ *B*|_*exp*_) are shown in Figure 2F

### Combining Tomo-seq with single-cell RNA-seq data

The TPM (transcripts per million)-normalized single cell RNA-seq (scRNA-seq) data collected from mouse olfactory epithelium available from (*54*) was used to identify cell-type specific genes. To this aim, we computed the average expression level for each cell type in the scRNA-seq dataset for all the differentially expressed genes that we identified in our TOMO-seq data. The genes with an average expression above 100 TPM in mOSNs and below 10 TPM in all other cell types were considered mOSN-specific. Conversely, genes with an average expression above 100 TPM in any of the non-mOSN cell types and below 10 in mOSNs were considered to be specific for non-mOSN cells.

### Gene Ontology (GO) enrichment analysis

GO Enrichment analyses were performed using the GOrilla online tool (http://cbl-gorilla.cs.technion.ac.il) with the option “Two unranked lists of genes (target and background lists)”. For each axis, we used as background list the list of the genes we tested.

### Identification of ligands and receptors associated with the NfiA, NfiB or NfiX transcription factors

The genes in the CellphoneDB ligands and receptor database (*87*) that were among our spatially differentially expressed genes were selected and Spearman correlation tests between their 1D expression patterns and the 1D patterns for the Nfi transcription factors were performed. Correlation coefficients from the three axes were averaged and FDRs from the 3 axes were combined with the Simes’ method (*112*) using the function combinePValues from the scran R library. Combined FDR values < 0.01 were taken as significant.

### 3D spatial reconstruction

The olfactory mucosa shape was obtained from publicly available images of the mouse nasal cavity along the posterior to the anterior axis published in (*66*). The area of the slices corresponding to the OM was manually selected and images of their silhouettes were made. Those images were then transformed into binary matrices having 1’s in the area occupied by the OM and 0’s in the remaining regions. The binary matrices were rescaled to match the spatial resolution in our dataset, which is composed of 54 voxels along the DV axis, 56 along the LML axis and 60 along the AP axis. Finally, matrices were piled in a 3D array in R to obtain an in-silico representation of the 3D shape of the OM, which, in total, was composed of 77,410 voxels. To perform the 3D reconstruction of the expression pattern for a given gene, first we normalized its expression levels by the volume of the slice at each corresponding position along the three axes, which was estimated using our 3D in silico representation of the OM. Then, we rescaled the data in such a way that the sum of the expression levels along each axis was equal to the average expression computed across the whole dataset. This rescaled dataset together with the binary matrix representing the 3D OM shape was used as input of the Iterative Proportional Fitting algorithm, which produced an estimation of the expression level of a gene in each voxel (*30*). Iterations stopped when the differences between the true and the reconstructed 1D values summed across the three axes was smaller than 1.

### Definition of zones by topic modelling

In order to identify zones, we fitted a Latent Dirichlet Allocation (LDA) (*115*) algorithm to the 3D gene expression patterns (in log10 scale) of the differentially expressed *Olfrs* (689 *Olfrs* x 77,410 voxels).

The LDA algorithm was originally employed for document classification: based on the words included in each document, LDA can identify “topics", in which the documents can then be classified. Using this linguistic analogy, in our application of LDA, we considered the genes as “documents”, and the spatial locations as “words”, with the matrix of gene expression levels being the analogous of the “bag-of-word” matrix (*70*). In this representation, the zones are the equivalent of “topics”, and they are automatically identified by LDA. We used the LDA implementation included in the R package “Countclust” (*116*), developed based on the “maptpx” library (*117*), which performs a maximum a posteriori estimation to for model fitting. LDA was run for all possible numbers of topics K ∈ [2,9]. The following parameters were chosen: convergence tolerance = 0.1; max time optimization step = 180 seconds; n_init = 3. For each number of topics k, three independent runs were performed with different starting points, in order to avoid biases due to the choice of the initial condition. We estimated the number of topics by computing the log-likelihood for each value of K ∈ [2,9]. As seen in Figure S3A, while the log-likelihood is a monotonically increasing function of the number of topic (as expected), for a number of topics around ~5 it shows a “knee” and starts to increase more slowly. This suggests that ~5 is the optimal number of topics needed to describe the complexity of the data without overfitting. Hence, we fix a number of topics equal to 5; however, we also verified that all our conclusions remain substantially unaffected if a different number of topics (e.g., 4 or 6) is chosen.

After running LDA with K=5, we retrieved the model output, which consists of two probability distributions: the first is P(position| k) with k ∈ [1,5], which is the conditional probability distribution defining the topic k; the second probability distribution is P(k | gene), namely the probability distribution that quantifies the degrees of belonging of a given gene to the topics k∈[1,5]. With these probability distributions, we can identify the spatial positions that form each topic and how the different topics can be combined to generate the spatial expression pattern of each gene.

Being a generative model, once trained, LDA can also decompose into topics the spatial expression patterns of genes that were not used during the training procedure. We exploited this feature of LDA to estimate the degrees of belonging of non-olfactory receptor genes. To this aim, we utilized an algorithm based on the python gensim library Lda.Model.inference function (*118*), using as input the estimated probability distribution P(position | k) with k ∈ [1,5]. The model fitting was performed using the Open Computing Cluster for Advanced data Manipulation (OCCAM), the High-Performance Computer designed and managed in collaboration between the University of Torino and the Torino division of the Istituto Nazionale di Fisica Nucleare (*119*).

### Definition of *Olfr* 3D indexes via diffusion pseudo-time

As explained in the section above, we can describe the spatial expression pattern of each gene through a set of five numbers, which represent the degrees of belonging to the five topics identified by LDA. We applied a diffusion map (*71*) to the degrees of belonging of the *Olfrs* to visualize them in two dimensions by using the “DiffusionMap” function from the “destiny” R package (*120*) (with distance=”rankcor” and default parameters). In this two-dimensional map, the *Olfrs* are approximately distributed along a curve that joins the most dorsal/medial genes (those in zones 1-2) with those that are more ventral/lateral (zones 3-5). To track the position of the genes along this curve, we computed a diffusion pseudo-time (DPT) coordinate (*72*) with the “DPT” function from the “destiny” R package (taking as starting point the gene with the smallest first diffusion component among the genes suggested by the function find_tips from the same package). In order to make the indexes go from Dorsal to Ventral, as in previous studies (*14*), we reversed the order of the DPT coordinates by substracting the maximum coordinate from all coordinates and multiplying them by (−1). By doing this, we obtained for each *Olfr* an index, which we called 3D index, representing its spatial expression pattern in the 3D space: more dorsal/medial genes (zones 1-2) have smaller 3D indexes than *Olfrs* expressed in the ventral/lateral regions (zones 3-5).

### Prediction of zone index for undetected *Olfrs* with Random Forest

We fitted a Random Forest model to the 3D indexes of 681 of the 689 *Olfrs* we characterized with our dataset (i.e., those that are located in genomic clusters). The following nine features of each *Olfr* were used as predictors: genomic position (i.e., gene starting position divided by chromosome length); genomic cluster; genomic cluster length; number of *Olfrs* in the genomic cluster; number of enhancers in the genomic cluster; cluster position (i.e., starting position of the cluster divided by the chromosome length); distance to the closest enhancer; gene position within the cluster (i.e., the distance of the gene starting position from the end of the cluster divided by the cluster length); and phylogenetic class.

These features were computed using the mm10 mouse genome in Biomart (*121*), while the list of enhancers and the genomic clusters assigned to each *Olfr* were taken from (*76*). The Random Forest model was fitted with the function “randomForest” (R library “randomForest” (*122*), with option “na.action = na.omit"). Afterwards, we performed a cross-validation test with the function “rf.crossValidation” from the “rfUtilities” package (*123*) with default parameters. Over 100 cross-validation iterations, the root mean square error (RMSE) was ≲10% of the mean 3D index. The feature importance was computed with the “importance” function from the randomForest library with default parameters. Finally, the Random Forest model trained on the 681 *Olfrs* was used to predict the 3D indexes of 697 *Olfrs* that were too lowly expressed or were undetected in our dataset. Overall, we were able to compute or predict with Random Forest a 3D index for all the *Olfrs* annotated in the mouse genome, except for 28 of them that do not have any genomic cluster assigned.

### Odorant information and Olfr-ligand pairs

All odorant structures and associated CAS numbers were retrieved from either Sigma-Aldrich (www.sigmaaldrich.com) or PubChem (https://pubchem.ncbi.nlm.nih.gov). A comprehensive catalog of the cognate mouse Olfr-ligand pairs was collected (last update: March 2021) by combining data from the ODORactor database (*124*) and additional literature sources (*4, 7, 100, 101, 125–151*).

This catalog includes 738 Olfr-ligand interactions for a total of 153 Olfrs and 221 odorants. These 153 *Olfrs* include 100 spatial *Olfrs* in our dataset and for which we have 3D indexes, and 49 additional *Olfrs* with predicted 3D indexes (see above). Next, we checked whether *Olfrs* pairs sharing at least one cognate ligand have more similar spatial expression patterns than pairs not sharing ligands. To do this, we computed the absolute values of the differences between the 3D indexes (Δ) of 1706 pairs of ORs sharing at least one odorant and 9922 pairs of ORs that are known to bind to different odorants (Figure 6B). The two sets of Δ values were significantly different (Mann-Whitney U test, p-value < 2.2×10^−16^).

### Correlation analysis of physico-chemical descriptors with 3D index

Physicochemical descriptors for ligands were obtained from the Dragon 6.0 software (http://www.talete.mi.it/). After removing the descriptors showing 0 variance, a table of 1911 descriptors for 205 ligands was obtained.

In addition to these, we estimated the air/mucus partition coefficients (*K_am_*) of the odorants as done previously (*21, 94*). Briefly, we calculated the air/water partition coefficients (*K_aw_*) for each odorant from the Henry’s Law constants obtained using the HENRYWIN model in the US EPA Estimation Program Interface (EPI) Suite (version 4.11; www.epa.gov/oppt/exposure/pubs/episuite.htm). Then, we computed the air/mucus partition coefficients (*K_am_*) according to the formula:

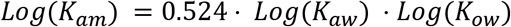

where *K_ow_* indicates the octanol/water partition coefficient, which were obtained using the KOWWIN model in the EPI Suite.

To increase the robustness of our correlation analysis, we removed the descriptors with 20 or more identical values across our set of ligands, and we initially considered only the ligands having 2 or more known cognate receptors; these filters gave us 1,210 descriptors (including K_am_) for 101 ligands.

We performed Spearman’s correlation tests between the physicochemical descriptors and mean 3D index of the cognate *Olfrs*, and we considered as statistically significant those correlations with FDR < 0.05 (see Supplementary Table S6). The correlations remained significant even when we removed *Olfrs* whose indexes were predicted with the Random Forest, or when only class II *Olfrs* were included in the analysis. In particular, the Spearman’s correlation of *K_am_* with the mean 3D indices is 0.69 (p-value = 2×10-3) when we removed *Olfrs* whose indexes were predicted with the Random Forest and 0.5 (p-value= 1.3×10-7) when only class II *Olfrs* were included.

### *In-situ* hybridization

*In-situ* hybridization was basically performed as previously described (Ibarra-Soria et al. 2017). 12-week-old male C57BL/6J mice were perfused with 4% paraformaldehyde and decalcified in RNase-free 0.45M EDTA solution (in 1x PBS) for two weeks.

Decalcified heads were cryoprotected in RNase-free 30% sucrose solution (1x PBS), dried, embedded in OCT embedding medium, and frozen at −80°C. Sequential 16 mm sections were prepared with a cryostat and the sections were hybridized to digoxigenin-labeled cRNA probes prepared from the different genes using the following oligonucleotides: Cytl (5’–AAAGACACTACCTCTGTTGCTGCTG – 3’ and 5’ –GTAAGCAGAGACCAGAAAGAAGAGTG – 3’), Moxd2 (5’ – TGTACCTTTCTCCCACTCCCTATTGTC – 3’ and 5’– CCCATGCAACTGGAGATGTTAATTCTG –3’), Olfr309 (5’–TACAATGGCCTATGACCGCTATGTG – 3’ and 5’– TCCTGACTGCATCTCTTTGTTCCTG – 3’), Spen (5’– GGTGGGAAACTTACCGGAGAACGTG – 3’ and 5’ – TGCTGCTGATGGAGTCACTACTG – 3’), Olfr727 (5’ – CGCTATGTTGCAATATGCAAGCCTC – 3’ and 5’– GCTTTGACATTGCTGCTTTCACCTC – 3’). The PCR products were cloned into pGEM-T Easy vector and the probes were obtained by in vitro transcription of the plasmids, using SP6 or T7 RNA Polymerases (Ambion) and DIG RNA Labeling mix (Roche).

## DATA AND CODE AVAILABILITY

RNA-seq raw data have been deposited and are publicly available as of the date of publication at ArrayExpress – accession number E-MTAB-10211. All original code and scripts for the 3D nose atlas shiny app has been deposited at Github and can be found [here](https://github.com/ScialdoneLab/Spatial-Transcriptomics/tree/master/atlas3Dnose). The 3D nose atlas processed data can be visualized and is publicly available here. Any additional information required to reanalyze the data reported in this paper is available from the lead contacts upon request.

## SUPPLEMENTARY MATERIAL

**Figure S1.**
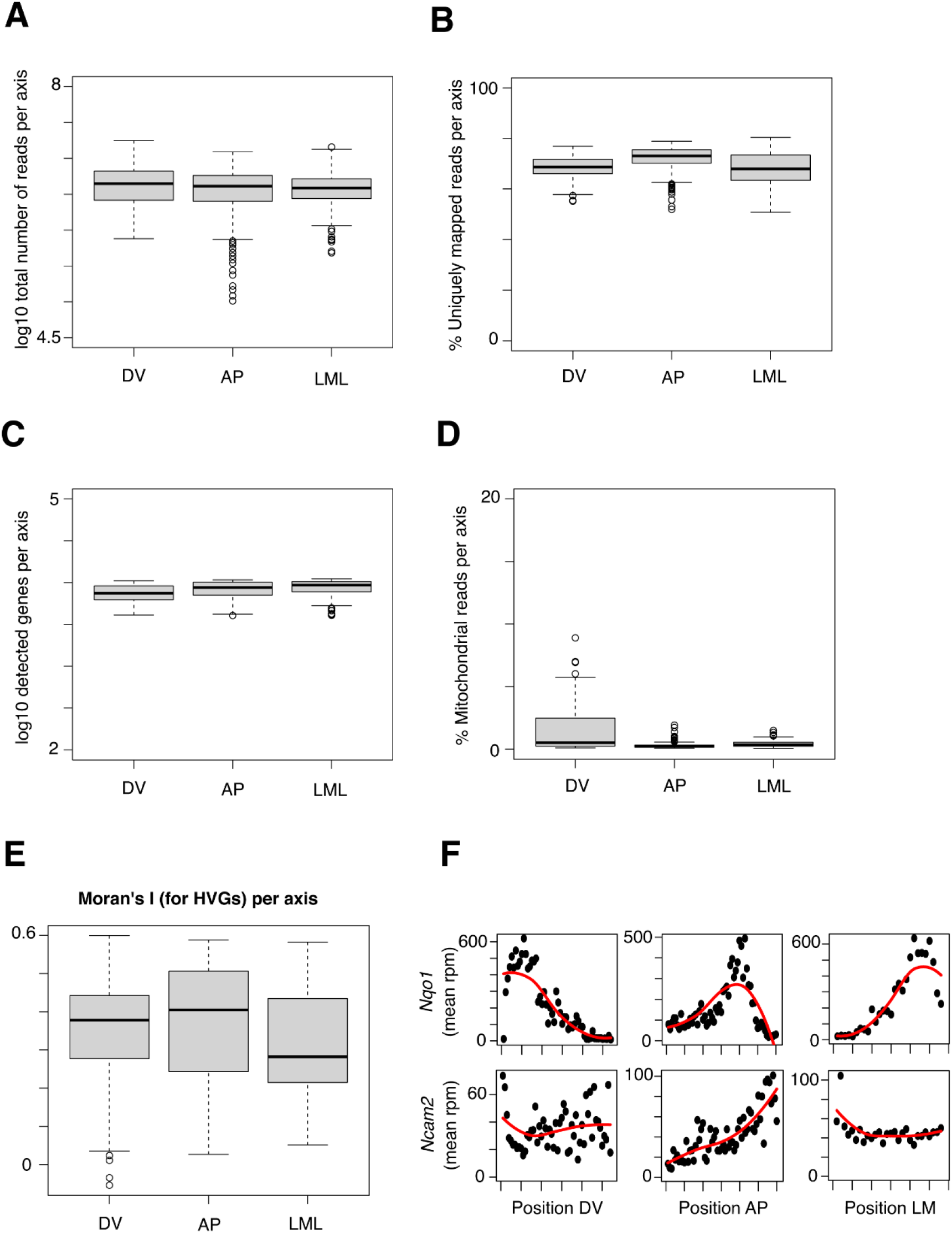
TOMO-seq data QC, related to Figure 1. (A) Boxplots showing the distributions of the log10 total number of reads per sample in each axis (DV = dorsal-ventral; AP = anterior – posterior; LML = lateral-mid-lateral). (B) Boxplots of percentage of uniquely mapped reads per sample per axis. (C) Boxplots of distributions of log10 detected genes per sample per axis. (D) Boxplots of percentage of mitochondrial reads per sample per axis. (E) Boxplots showing the distribution of the Moran’s I statistics calculated for the top 100 Highly Variable Genes per axis. P-values are computed for each gene and then combined with the Simes’ method. The combined p-values are < 2.2×10^−16^ for all axes. (F) Normalized expression of canonical OM spatial marker genes along the 3 axes. Red line showing fits with local polynomial models.

**Figure S2.**
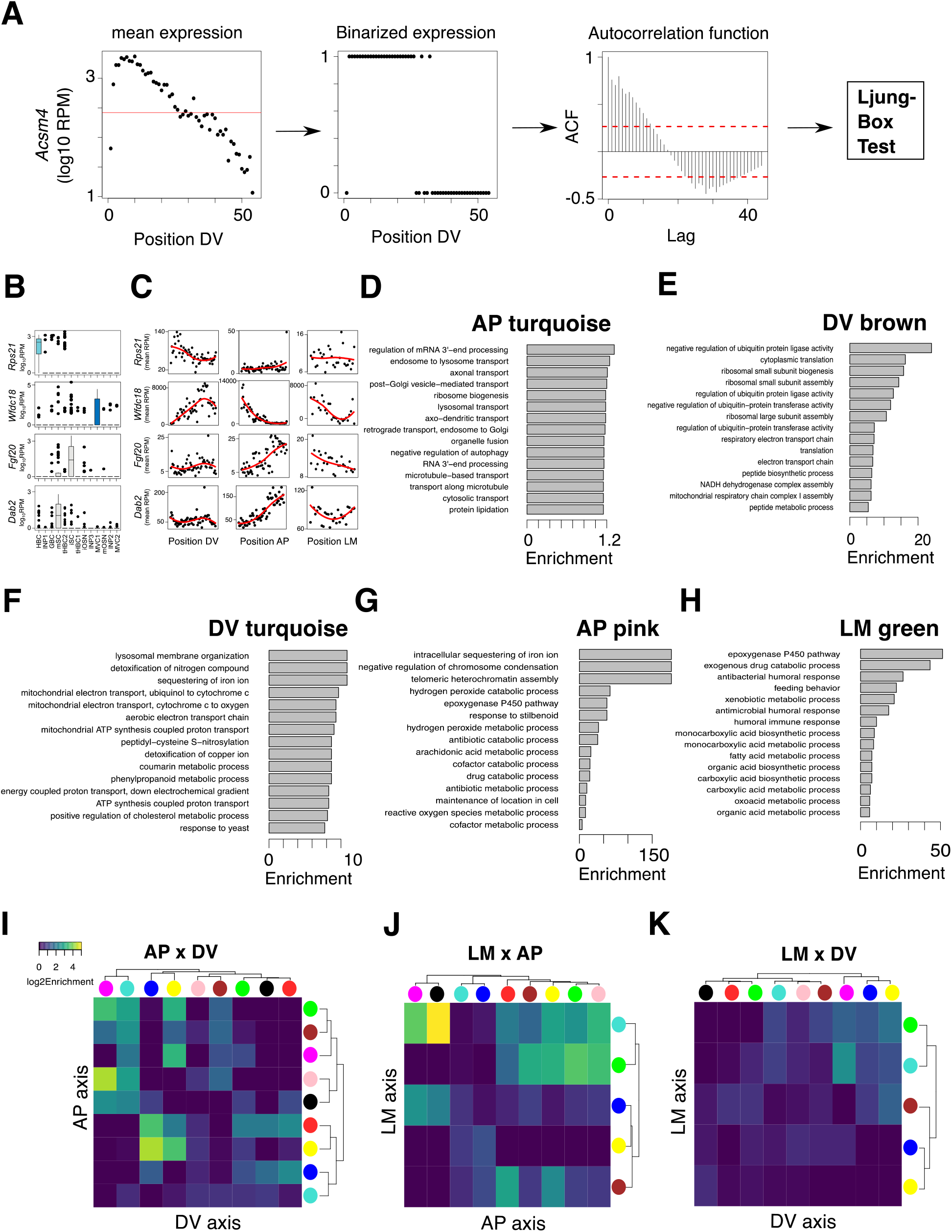
Spatial differential expression analysis, related to Figure 2. (A)Schematics of strategy to find spatially differentially expressed genes; as an example, data for Acsm4 along the dorsal-ventral (DV) axis is shown: Gene expression was binarized according to whether the expression per slice was higher or lower than the median expression (red horizontal line). Then, we computed the autocorrelation function for different values of the lags, and we applied the Ljung-Box test to verify whether the autocorrelation values are significantly higher than zero. (B) Box plots of example genes’ expression (log10 reads-per-million, RPMs) distributions in different cell types. None of these genes is expressed in mOSNs (INP = Immediate Neuronal Precursors; GBC = Globose Basal Cells; mOSNs = mature Olfactory sensory neurons; iOSNs = immature Olfactory Sensory Neurons; MVC = Microvillous Cells; iSC = Immature Sustentacular Cells; mSC = Mature Sustentacular Cells; HBCs = Horizontal Basal Cells). (C) Spatial gene expression trends along each axis of the example genes shown in panel B. (D-H) Bar plots showing the enrichment values for the top enriched Gene Ontology categories in different gene clusters (see Figure 2E). (I-K) Heatmap showing the log2 enrichment for the intersection between different gene clusters (indicated by colored circles) across pairs of axes, after excluding *Olfr* genes.

**Figure S3.**
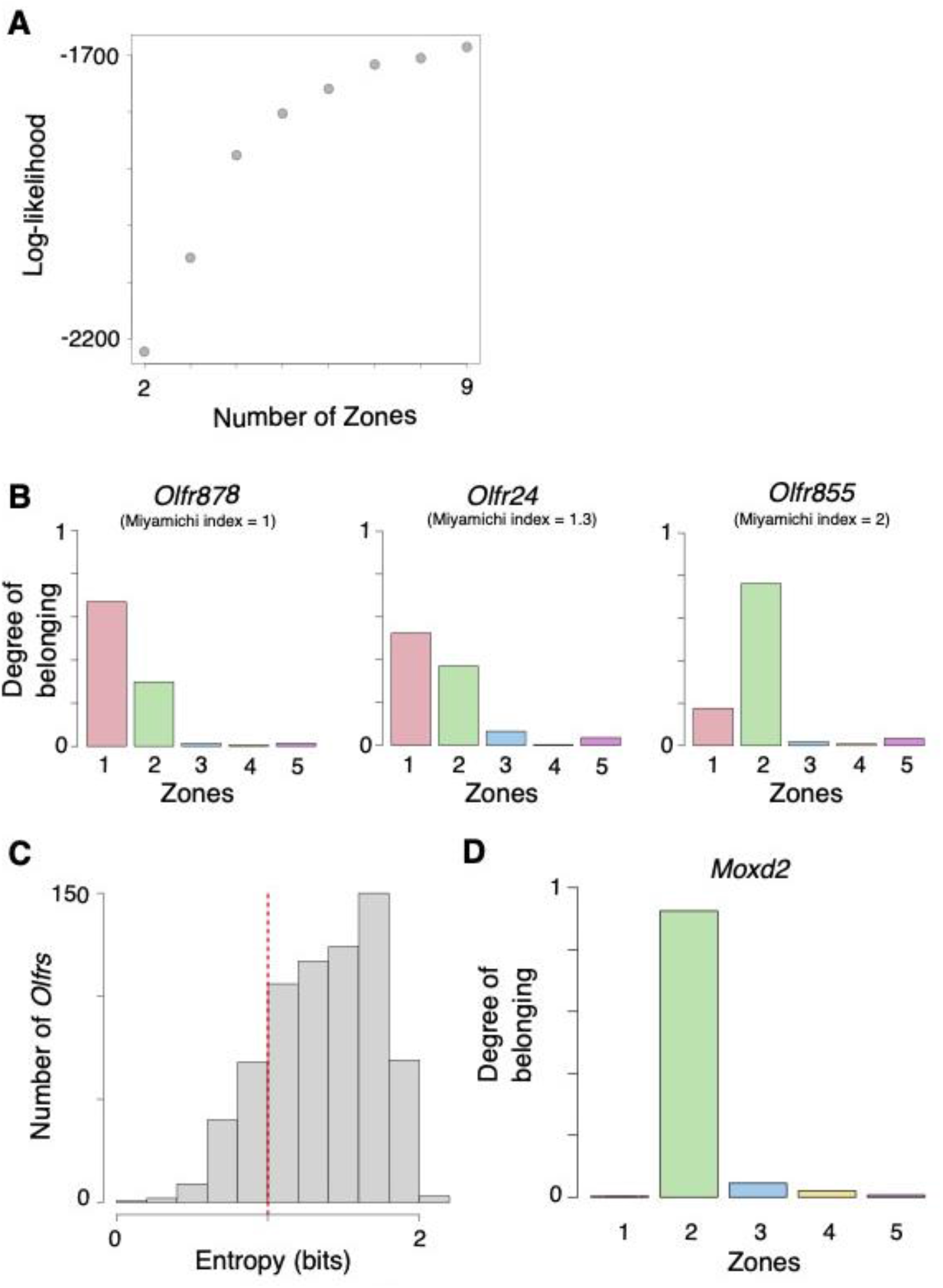
Olfr genes 3D zones, related to Figure 3. (A) Log-likelihood values for fits with LDA models as a function of the number of zones. (B) Bar plot showing the degrees of belonging of *Olfr* genes with overlapping spatial patterns (Miyamichi indexes of 1, 1.3 and 2 respectively). (C) Distribution of entropy values of our 689 Spatially differentially expressed *Olfrs’* spatial expression. The *Olfrs* with entropy values less than 1 bit (vertical red line) were considered to fit mostly in one zone. (D) Bar plot showing the degrees of belonging of *Moxd2*.

**Figure S4.**
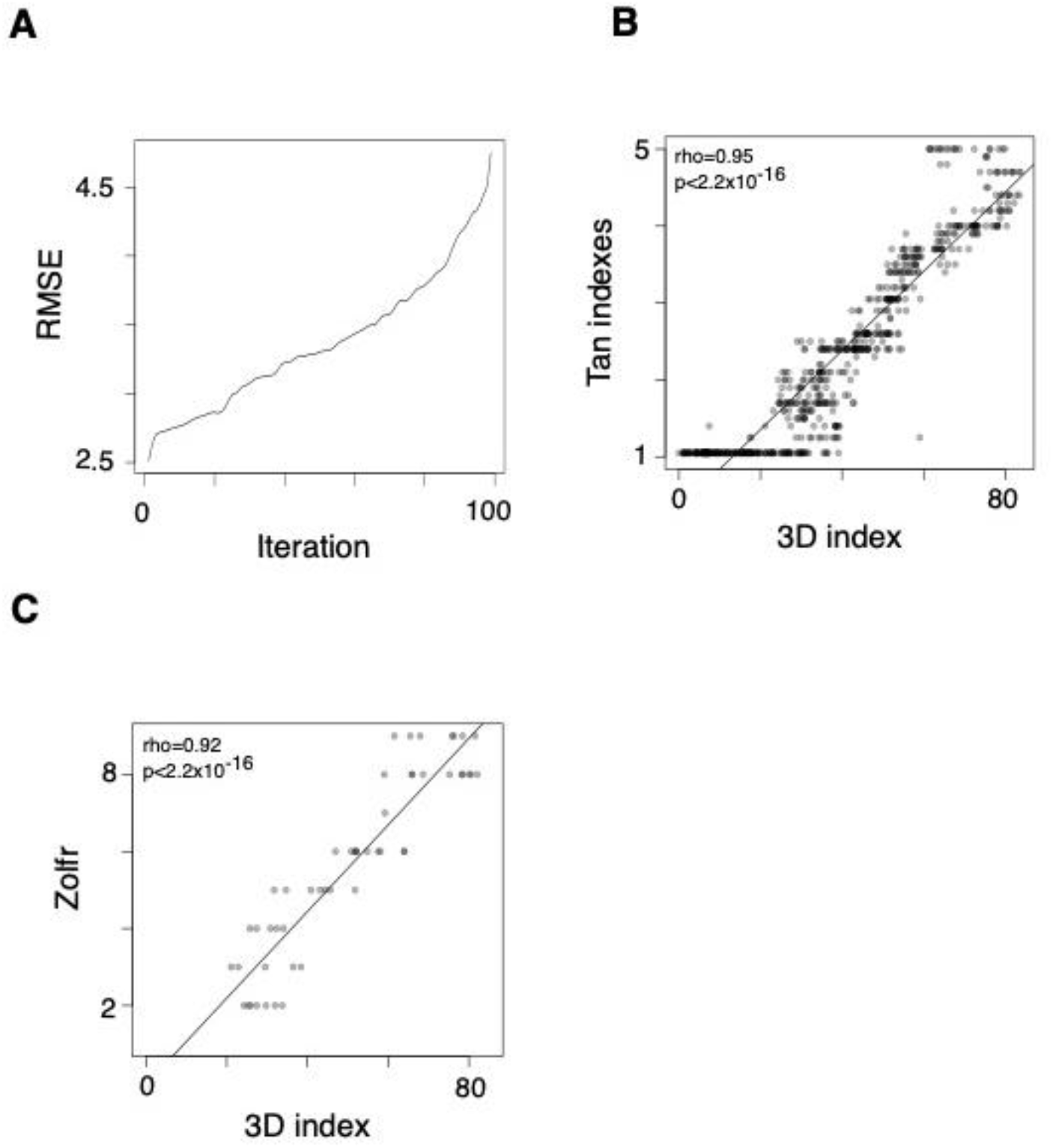
Olfr 3D index prediction, related to Figure 4. (A) Root mean square error (RMSE) per iteration of the cross-validation test for the Random Forest model used to predict 3D indexes. (B) Scatter plot showing the correlation of our 3D indexes with the zone indexes estimated by [66] (“Tan Indexes”), who performed RNA-seq on 12 samples at different positions along the dorsal-ventral axis of the OM and estimated indexes using as reference the ~80 *Olfrs* analyzed in [14] via ISH. (C) Scatter plot illustrating the comparison of our 3D indexes versus the zones defined by [16] (“Zolfr”) from ISH data. For this comparison, these zones were numbered from 1 to 9 from the most dorsal to the most ventral.

**Figure S5.**
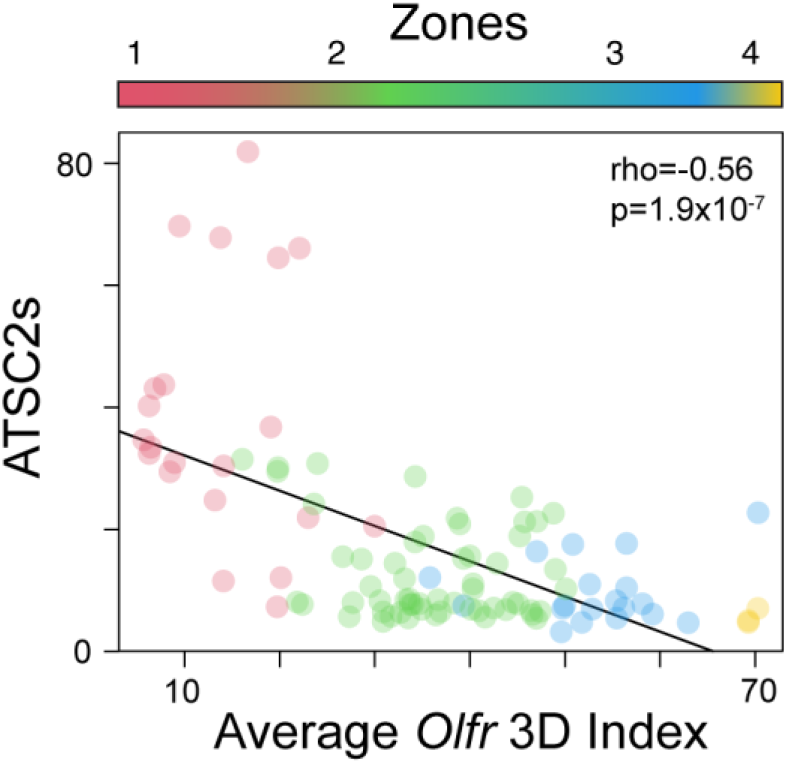
Physiological role of the zones, related to Figure 6. Scatter plot illustrating the correlation between ATSC2s of the odorants and the average 3D indexes of their cognate Olfrs. Only odorants for which we know at least two cognate Olfrs (110) were used here. Odorants are colored according to the zone they belong to (defined as the zone with the highest average degree of belonging computed over all cognate receptors).

---

**Table S1. Microsoft Excel format file, related to Figure 1. Quality control.** Sheets 1-3: Quality statistics of TOMO-seq data from each sample (one per axis). Sheet 4: Number of detected genes per axis (Genes were considered as detected when they had at least one (RPM) count in at least 10% of the samples from one axis).

**Table S2. Microsoft Excel format file, related to Figure 2. Gene expression in different cell types.** Sheet 1: Spatially differentially expressed Genes not expressed in mature olfactory sensory neurons per axis. Sheet 2: Gene Ontology Enrichment analysis results (output from GOrilla) for the spatially differentially expressed genes not expressed in mature olfactory sensory neurons. Sheet 3: Total number of spatially differentially expressed Genes (totalDEGs), number of spatially differentially expressed genes coming from mature olfactory sensory neurons (OSNsDEGs), and number of spatially differentially expressed genes not expressed in mature olfactory sensory neurons (nonOSNsDEGs).

**Table S3. Microsoft Excel format file, related to Figure 4. Spatially differentially expressed genes.** Sheets 1-3: genomic features (Ensembl ID, gene name, location and length), analysis-related features (p-value, FDR, and cluster) of spatially differentially expressed genes (one table per axis).Sheets 4-23: Gene Ontology Enrichment analysis results (output from GOrilla) for the spatially differentially expressed genes per cluster.

**Table S4. Microsoft Excel format file, related to Figure 4. 3D indexes.** Sheet 1: Log likelihood values for LDA topic models with numbers of topics from 2 to 9. Sheet 2: Degrees of Belonging, zone with maximum DoB, entropy and 3D indexes for spatially differentially expressed Olfr genes.Sheet 3: Genomic features (genomic cluster, length of genomic cluster, number of Olfr genes in the genomic cluster, number of enhancers in the genomic cluster, distance from gene to closest enhancer, Olfr class, gene chromosomal position, gene position in the genomic cluster, and cluster chromosomal position) and predicted 3D indexes for Olfr genes with available data.

**Table S5. Microsoft Excel format file, related to Figure 5. Non-Olfr genes.**

**Table S6. Microsoft Excel format file, related to Figure 6. Olfr-ligand analysis related data.** Sheet 1: List of Olfr – ligand pairs. The organoleptic properties of some odourants and the references are also listed. Sheet 2: Log (air / mucus partition coefficient) for ligands with available information. CAS numbers and SMILES included.

